# Neural Ordinary Differential Equations Inspired Parameterization of Kinetic Models

**DOI:** 10.1101/2024.12.20.629595

**Authors:** Paul van Lent, Olga Bunkova, Lèon Planken, Joep Schmitz, Thomas Abeel

**Affiliations:** Delft Bioinformatics Lab, Delft University of Technology, Delft, Zuid-Holland, Netherlands; Department of Science and Research, dsm-firmenich, Delft, Zuid-Holland, Netherlands; Infectious Disease and Microbiome Program, Broad Institute of MIT and Harvard, Cambridge, MA, United States of America

**Author notes:** **Author contributions**: Paul van Lent (Conceptualization, Methodology, Software, Writing - original draft preparation), Olga Bunkova (Conceptualization, Methodology, Software, Writing - Review & Editing), Lèon Planken (Software, Writing - Review & Editing), Joep Schmitz (Conceptualization, Supervision, Writing - Review & Editing), Thomas Abeel (Conceptualization, Supervision, Writing - Review & Editing). Delft Bioinformatics Lab, Delft University of Technology, Delft, Zuid-Holland, Netherlands.

## Abstract

**Motivation:** Metabolic kinetic models are widely used to model biological systems. Despite their widespread use, it remains challenging to parameterize these Ordinary Differential Equations (ODE) for large scale kinetic models. Recent work on neural ODEs has shown the potential for modeling time-series data using neural networks, and many methodological developments in this field can similarly be applied to kinetic models.

**Results:** We have implemented a simulation and training framework for Systems Biology Markup Language (SBML) models using JAX/Diffrax, which we named *jaxkineticmodel*. JAX allows for automatic differentiation and just-in-time compilation capabilities to speed up the parameterization of kinetic models. We show the robust capabilities of training kinetic models using this framework on a large collection of SBML models with different degrees of prior information on parameter initialization. Finally, we showcase the training framework implementation on a complex model of glycolysis. These results show that our framework can be used to fit large metabolic kinetic models efficiently and provides a strong platform for modeling biological systems.

**Implementation:** Implementation of *jaxkineticmodel* is available as a Python package at https://github.com/AbeelLab/jaxkineticmodel.

**Author summary:** Understanding how metabolism works from a systems perspective is important for many biotechnological applications. Metabolic kinetic models help in achieving understanding, but there construction and parametrization has proven to be complex, especially for larger metabolic networks. Recent success in the field of neural ordinary differential equations in combination with other mathematical/computational techniques may help in tackling this issue for training kinetic models. We have implemented a Python package named *jaxkineticmodel* that can be used to build, simulate and train kinetic models, as well as compatibility with the Systems Biology Markup Language. This framework allows for efficient training of kinetic models on time-series concentration data using a neural ordinary differential equation inspired approach. We show the convergence properties on a large collection of SBML models, as well as experimental data. This shows a robust training process for models with hundreds of parameters, indicating that it can be used for large-scale kinetic model training.

## Introduction

Kinetic modeling is a useful tool for describing biological systems in a quantitative manner, with many applications in the biotechnological and medical domain [1, 2]. In the biotechnological domain, the application of kinetic models includes assessing metabolic control of pathways [3], simulation of metabolic engineering scenarios [4, 5], and optimizing feeding strategies on the bioprocess level [6]. Effective deployment of kinetic models for these purposes requires a representative description of the biological process in mathematical equations, as well as fitting the model parameters to available data. The encountered data in this domain is typically limited and either steady-state or dynamic in nature.

Metabolic kinetic models are described by ODEs that describe the change over time of metabolites (m) by a right-hand-side formula that consists of the mass balances imposed by the stoichiometric matrix (S) and a vector of reaction flux functions (v⃗) (eq. 1). The process of finding a model that reproduces observed data therefore consists of establishing the stoichiometric matrix [7, 8], determining kinetic mechanisms [9], and parametrization of the flux functions.

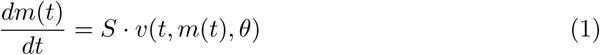

Many parameter estimation methods have been proposed and implemented in publicly available software packages. Some toolboxes focus specifically on fitting steady-state data [10–12], while other toolboxes provide more general purpose parameter inference methods for metabolic modeling, such as pyPESTO [13]. pyPESTO provides an interface to many different optimization schemes: local versus global, gradient-free versus gradient-based optimizers [13]. Particularly efficient optimization methods are the local gradient-based methods, with the downside that the chance of finding a local optimum is higher compared to global optimization methods.

Fitting parameters to large-scale kinetic models can be a difficult task [14]. Since biology operates on many different timescales, the scale of biologically inspired parameters can vary by orders of magnitude [15]. Additionally, the output of ODEs might be insensitive to many parameters, sometimes referred to as sloppiness [16]. These characteristics of the parameter space complicate the use of (gradient-based) local optimization schemes. Furthermore, many kinetic models in systems biology have unidentifiable parameters, which complicates the fitting process even more [17]. Finally, most biological systems are known to be stiff, which leads to problems when numerically solving the ODEs [18].

Recently, Neural ODEs were introduced [19] and applied to modeling time-series concentration data [20]. The idea behind a neural ODE is to replace the right-hand-side of equation 1 by a neural network and then use a numerical solver to predict a time-series. The neural network is then trained using back-propagation with the adjoint state method; a very efficient method for estimating gradients required in stochastic gradient descent (see Fig. S1.) [19]. Even though neural ODEs lack the necessary mechanistic structure required in biotechnological/medical applications, many of the techniques for training, such as adjoint gradient computation, can similarly be applied to metabolic kinetic models. This paves the way to fitting large-scale kinetic models.

In this work, we have implemented a *JAX* -based systems biology training framework using *Diffrax* [21, 22], which we named *jaxkineticmodel*. The training framework is tailored to systems biology models and is compatible with the Systems Biology Markup Language [23]. Training is performed by gradient descent in log parameter space [15] with a stiff numerical solver [24] and a custom loss function to deal with metabolic scale differences. To further stabilize training, we perform gradient clipping, a method that is widely used in stabilizing the training process of recurrent neural networks [25]. We apply this implementation on a large collection of SBML models to answer questions on robustness of the training procedure in terms of convergence properties, as well as a post-hoc analysis of the parameter space. Finally, we show the training of a large-scale kinetic model of glycolysis (141 parameters) to model feast/famine feeding strategy datasets [26].

## Results

### Training SBML models using *Diffrax*

In order to analyze the behavior of the parameterization using Neural ODEs, an efficient and easy-to-use ODE simulation tool for systems biology models with automatic differentiation options for calculating adjoint sensitivities was required. We have implemented a *JAX* -based simulation tool and training framework that is compatible with Systems Biology Markup Language (SBML) models in *Diffrax* [21, 22] (Fig. 1). SBML models are the standard accepted format for saving systems biology models in a reproducible manner [23].

**Fig 1.**
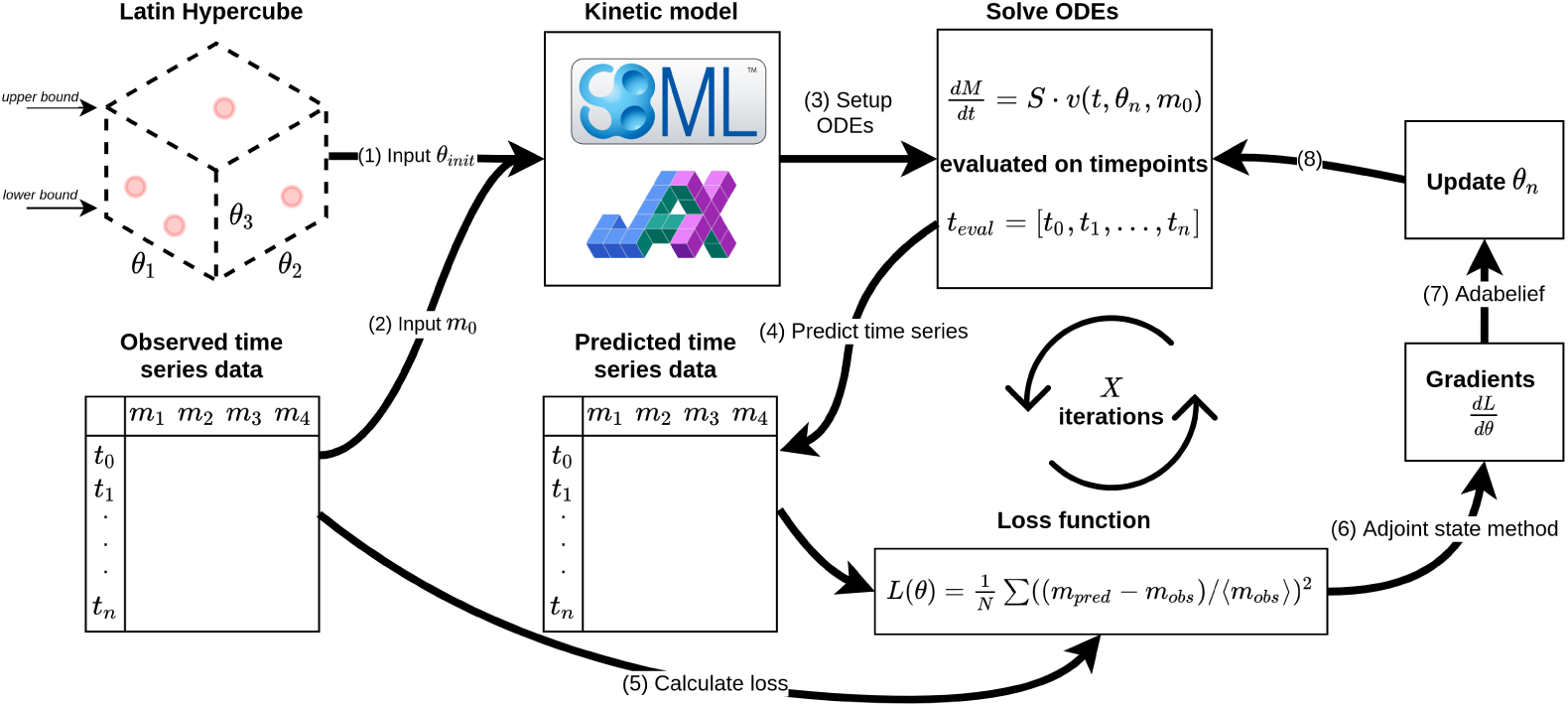
Overview of the implemented simulation tool and training framework in *Diffrax*. A tool for converting SBML models to a *JAX* -compatible model that can be simulated using *Diffrax* was implemented [21, 22]. Latin Hypercube sampling is used to initialize parameters given a lower- and upperbound value (1). The initial conditions are retrieved from the observed dataset (2) and are used to setup the ODEs for simulation (3). After predicting a time-series dataset given the initial guess (4), the mean-centered loss is calculated (5). The gradients that are calculated through the adjoint state method (6) are then used to update parameters using AdaBelief (7) [54]. This process is repeated for X steps or until convergence (8).

The training input consists of three information modes. The kinetic model is loaded from an SBML format or a manually implemented *JAX* -compatible class that allows for Just-In-Time (JIT) compiling. The observed time-series data that is used for fitting is used to get the initial conditions from t_0_. Finally, an initial parameter guess is required, which was obtained using Latin Hypercube Sampling [56]. Due to nonlinearities and potential non-convexity of the solution space, multiple initialization are typically required.

For the training process, the ODEs are solved for timepoints that are observed in the dataset and the loss function is calculated. Due to large differences in metabolite concentration ranges, a mean-centered loss function was used to ensure roughly equal contribution of metabolites to the mean squared error. Finally, N iterations of stochastic gradient descent using AdaBelief can be performed [54].

While kinetic models typically have a more mechanistic structure of the right-hand-side of dm(t)/dt, similar tools that are used to train Neural ODEs (e.g., the adjoint state method) can be applied. The goal of the study performed here is to address properties on the convergence of local gradient-based methods for a large collection of systems biology models.

### Convergence analysis reveals robust training of systems biology models

Training kinetic models using the Neural ODE framework requires an initial guess of parameters. This requires setting a lower- and upper bound of parameters for the initialization and sampling the space using any sampling method. Due to the dependency of the loss function on numerical integration of the ODEs, not every parameter initialization might be successful. This can be attributed to either stiffness of the dynamical system or unstable behavior of the systems given that particular initial guess [18]. We therefore research the performance in terms of initialization success and the training process for different priors and models of varying sizes.

To motivate the main analysis of SBML models, we show the influence of parameter bounds on one model when sampling 100 initializations using Latin Hypercube Sampling (Fig. 2A) [56]. The percentage of models that are below the loss threshold are reported for five different parameter bound priors. It is observed that with larger bounds, the percentage of successfully trained models decreases, as would be expected. We also compare the convergence success among five different systems biology models with a fixed prior bound ( <u>^1^</u> θ*_true_* ≤ θ*_true_* ≤ 10θ*_true_*) (Fig. 2B). This allows for comparing the performance of the Neural ODE framework for parametrization across a large collection of Systems Biology models. In order to compare many models across different bounds, we chose a loss threshold of 10*^−^*^3^.

**Fig 2.**
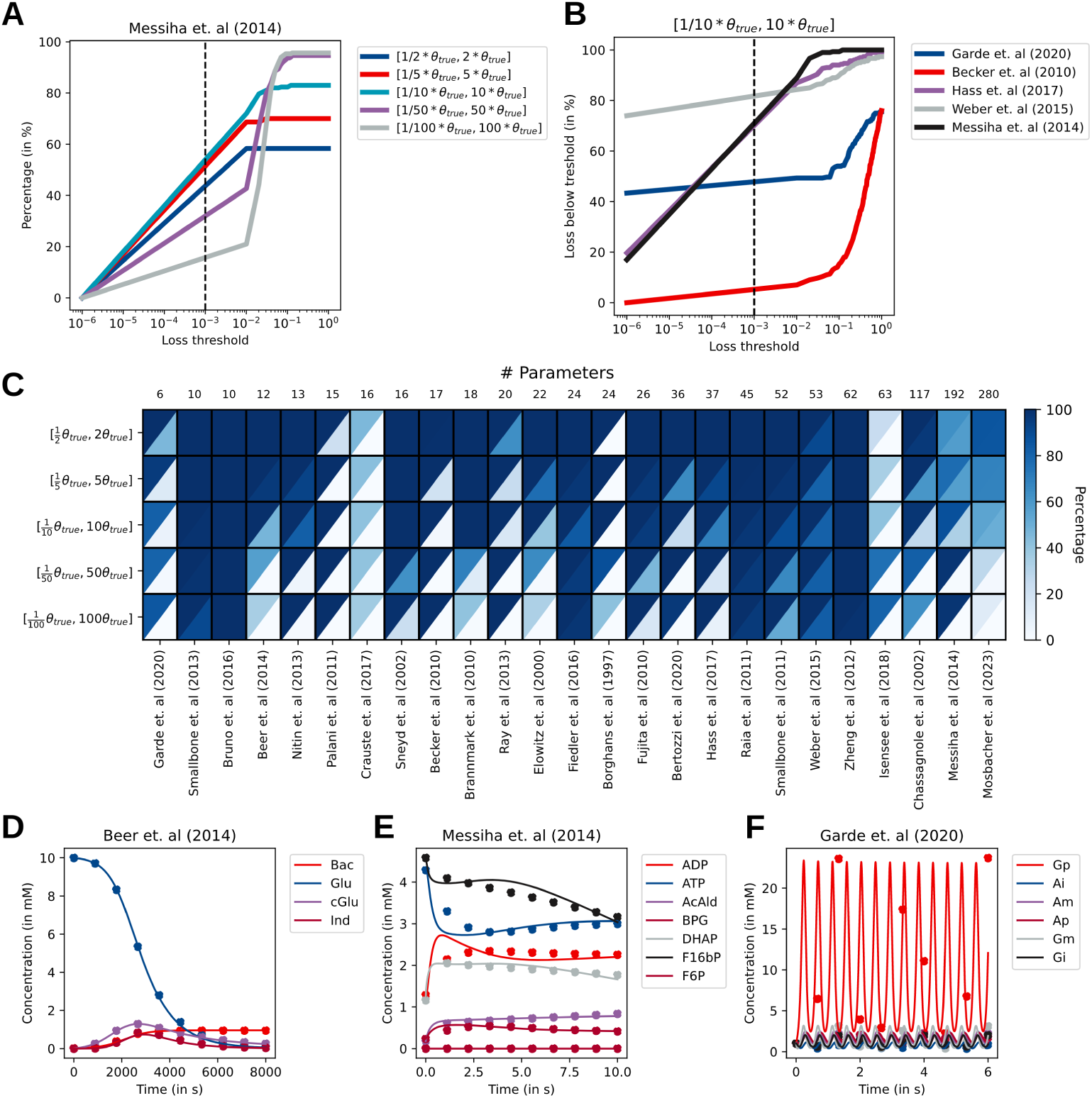
Analysis of convergence properties for a collection of SBML models. Latin hypercube sampling is performed for five different lower and upper bound priors of the parameters and training is performed. A) Percentage of successful convergence of an example model, where success is defined as the percentage of models below the loss threshold on the x-axis. B) A similar plot, but now with a fixed prior and five SBML models. C) Heatmap for the initialization success percentage (upper diagonal) and training success (lower diagonal) for *loss* < 10*^−^*^3^. D,E,F) Examples of fitted SBML models after training.

### Initialization success and the training process is stable across models and priors

Figure 2C shows a heatmap for 25 SBML models for five different priors. The left column of each model is the initialization success percentage, while the right column is the percentage of models after training that were below the loss threshold (10*^−^*^3^).

Overall, we see a high initialization success percentage for most SBML models. For many models, the loss function can be calculated independent of the bounds used in this study. For six models, the behavior of decreasing initialization success given the priors is observed, in line with what would be expected [31, 37, 39, 40, 50, 52].

Interestingly, a reversed pattern is observed for two models, where the initialization success increases with larger bounds [49, 51]. For one model, the initialization success is low, independent of the prior [34].

The training process consists of 3000 iterations of stochastic gradient descent using AdaBelief and a global norm clipping with a learning rate of 10*^−^*^3^ [25, 54]. The right column of each model for Figure 2C shows the percentage of successfully trained parameter given the initialization success. Here, the effect of priors is observed more clearly. For the smallest bounds, we observe good convergence to the global minimum for most models, although not all models are successfully trained. This can be in some cases because the loss is already close to the threshold and only requires a few steps of gradient descent, but often the chance of finding the right parameter set is also important from a perspective of stability and stiffness. Furthermore, it is observed that many models have a decreasing percentage of successfully trained initialization with increasing bounds. Generally speaking, we do not observe a clear relation between the number of parameters (or state variables) and the difficulty of training these models, except that the computation time increases for larger scale models.

To further observe whether the models fit data such that it properly captures the metabolite concentration dynamics, we simulate three models with losses below the threshold given in the heatmap for their best fit parameters with bounds 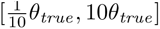 and compare it to the simulated data points (Fig. 2D-F). It can be observed that for the model from Beer et. al (2014) and Messiha et. al (2014) the dynamics are fairly close if not perfectly matching the simulated data points (Fig. 2D,E). For the model from Garde et. al (2020), despite the loss being below the threshold, the trained dynamics are not close to the true dynamics, but rather finds a heavily oscillating parameter set that then matches the data. This behavior is not observed when the prior is between one-fifth and five of the true parameters. One way to mitigate this when a good prior is not available could be to filter out parameter sets that have periodicity that lead to unexpected periodicity. Furthermore, increasing the number of data points to train on might also decrease the likeliness of finding a local optimum.

Overall, initialization success is stable across a wide variety of models, which increases the likelihood of effectively learning parameters. While we see for some models a dependency on the bounds, this effect can be mitigated by increasing the initial sampling size. The loss after training shows a clearer effect of the priors. This indicates that when parameterizing kinetic models, the prior can be of high importance.

Although it might in practice be difficult to define strict bounds for parameters *a priori*, using kinetic databases like Brenda might be a way to set an initial parameter. [16, 58].

### Small subset of parameters explain dynamic behavior in systems biology models

As it was established that kinetic models could be trained using the Neural ODE framework, we sought to explain the importance of parameters during training, as it has been reported that many parameters in systems biology models are *sloppy* : that is, parameters for which the model is extremely insensitive to the actual value [16]. These parameters are thought to complicate the training process, but also give an indication about the parameters that matter for the observed dynamic behavior of the system.

One of the SBML models that were retrieved from a previous collection of benchmark model was used for further analysis [15, 36]. The model describes the dynamic behavior of the epoR system, which is a widely studied pathway. We start the analysis by computing the cosine distance (see *Methods*) between the true parameter set from the SBML model and the initialized parameter set as well as the trained parameters (Fig. 3A). There seems to be no clear difference in the average distance before and after training, although a small increase in density is observed closer to the optimum for the trained parameters. This suggests that the average distance is not directly influenced during training. However, when performing Principal Component Analysis on the parameters (see *Methods*), a clear separation is observed between the parameter before -and after training (Fig. 3B). The trained parameters are closer to the true optimum in parameter space, suggesting that training has in fact been successful. When looking at the loadings of the PCA, we observe that for the first principal component a few parameters are of high importance for this separation (Fig. 3C). Indeed, when we take only the first three parameters and redo the cosine distance plot, we observe that these parameters are very close to the true optimum, suggesting that only a subset of parameters are important to be precisely set (Fig. 3D). This is in the literature known as *sloppy* models and has been suggested to be a general feature in systems biology models, either through the way they are constructed, or due to the actual underlying biological properties of robustness to parameters [15, 16]. We observe the same behavior for other SBML models, suggesting that this is a general feature of this type of systems. Furthermore, we observe that the parameters are typically distributed in log-space or even on an exponential distribution. Alltogether, these results indicate that the training process can also be used to identify the importance of parameters by looking at the behavior of the ensemble of parameter initialization after training.

**Fig 3.**
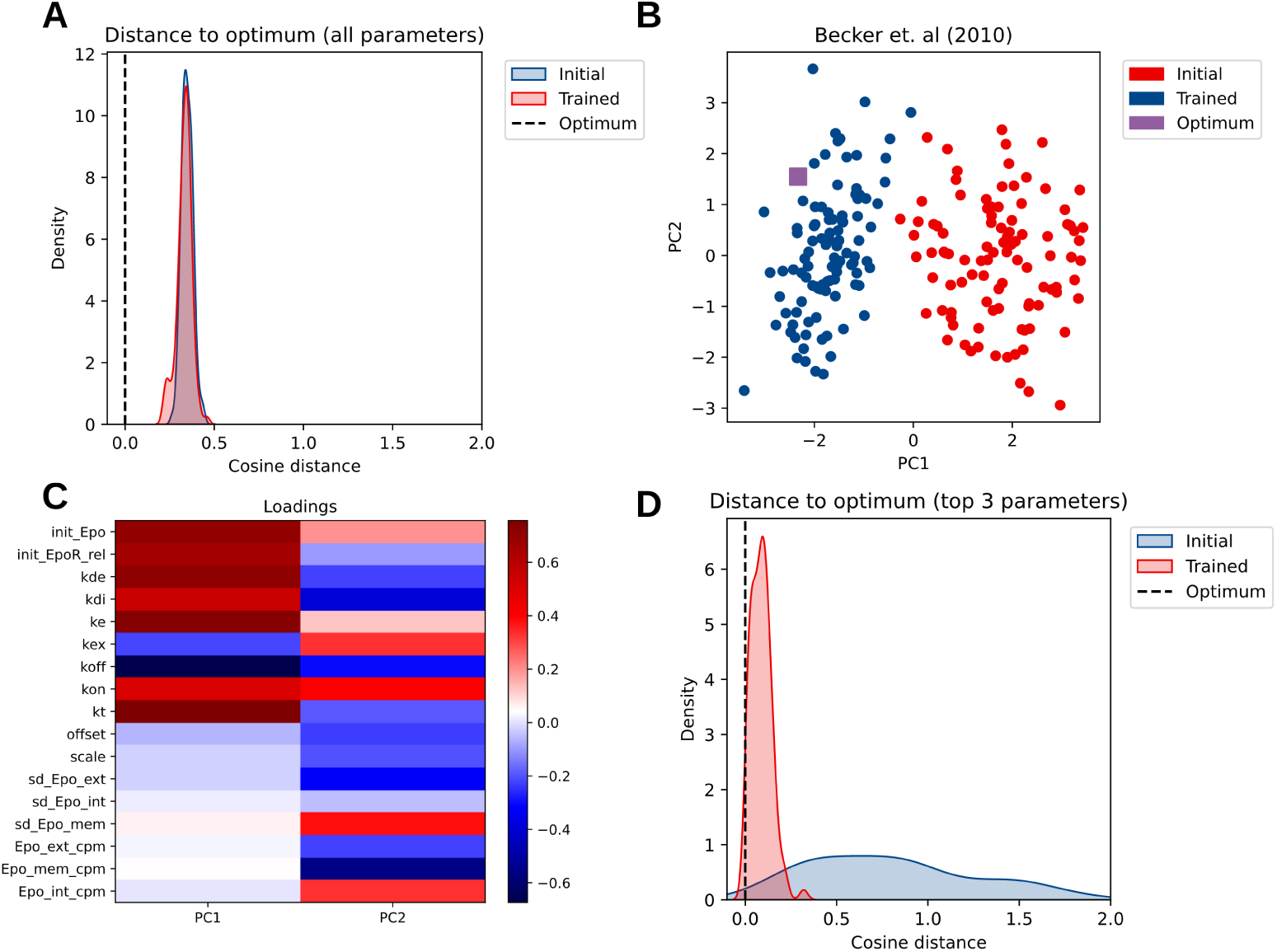
Only few parameters are precisely set during training. A) The cosine distance of the initialization parameter set and trained parameter set are compared to the true optimum parameter set. B) Principal Component Analysis of the initialization parameter set and parameters after training. The trained parameter lie closer in space to the optimum, although much variation exists. C) Loadings plot of the first two Principal Components reveal some parameters with large impact on covariance. D) Cosine distance for the top three parameters with the highest loadings. The distance is closer to the optimum, indicating a precise setting of some parameters.

### Large scale kinetic models can be trained using the neural differential equation framework: an example of glycolysis

While we have shown that neural differential equations could be used for a large variety of different SBML models, real time-series metabolomics data has additional complexity in terms of heterogeneity of the measured metabolites as well as noise. We therefore reimplemented a kinetic model of glycolysis in *JAX* [22, 57]. This model consists of 20 metabolites, 37 reactions, and 141 parameters. The kinetic mechanisms used are implemented as *Jax* compatible classes for well-known equations (e.g., Michaelis Menten) that can be reused for other purposes (see Table SI1).

The model was initialized with literature values reported in the previous MATLAB implementation [57]. 10000 iterations of stochastic gradient descent using AdaBelief was performed, resulting in the fit as reported in Figure 4. The panels show the dynamic response of a glucose pulse during a feast-famine cycle for some previously reported datasets [26]. The striped lines are inferred dynamic responses of metabolites, for which no training data was available. The glucose pulse obviously leads to an increase through the upper- and lower glycolysis, as well as a pulse through the glycerophospholipid pathway. In the first steps ATP is formed, which is well-captured by the dynamic response of the model. There is also a drop in NAD+ due to an increased activity through the lower glycolysis, where NAD+ is consumed by *glyceraldehyde-3-phosphate dehydrogenase*.

**Fig 4.**
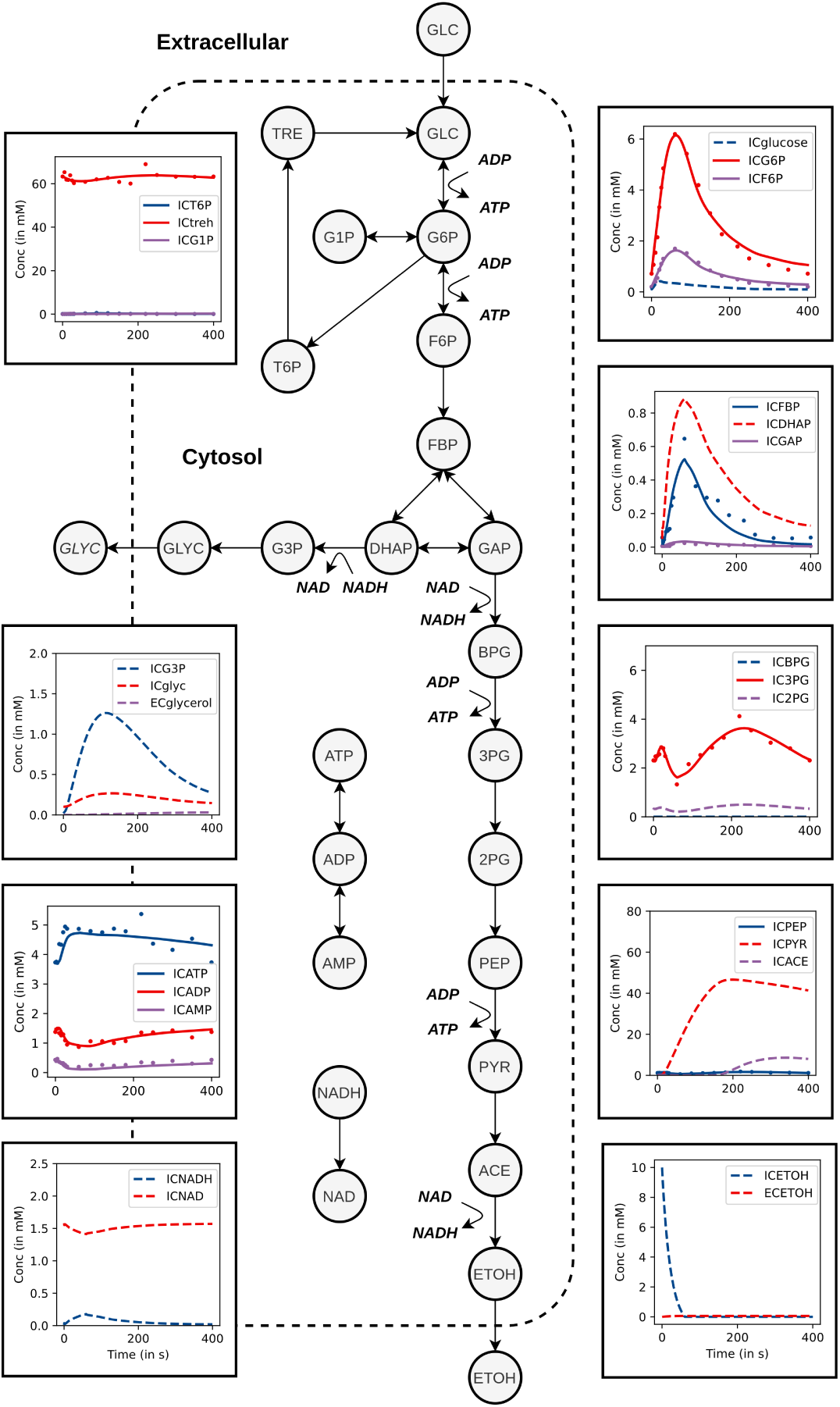
Fitting a feast/famine cycle to a glycolysis model. A simplified schematic of glycolysis, along with panels of the dynamic response of glycolysis to a glucose pulse. Dots are the measured metabolites over time, while the lines indicate the prediction by the model. Dotted lines are inferred metabolite dynamics, where there was no available data.

Overall, a good fit is observed between modeled and measured data. Even though this is a relatively large and complicated model of glycolysis, the fitting process for one dataset takes only a few hours. This shows that the implementation could be used to fit large-scale kinetic models. Furthermore, the training process is easy to extend to fit multiple datasets simultaneously (see Fig. S4).

## Discussion

Large-scale kinetic models have the potential to explain systems level behavior of biological systems, but their parametrization has proven to be difficult [14]. In this work, we have implemented a neural ODE inspired training framework tailored to systems biology models. Due to the *JAX* -based simulation software *Diffrax* training is relatively fast, which paves the way to large-scale kinetic model training [21, 22]. The implementation provided in this study is publicly available at https://github.com/AbeelLab/jaxkineticmodel.

While in original neural ODEs the right-hand-side of equation 1 is replaced by a neural network, we maintain the rigid structure of metabolic kinetic models for two reasons. First, the amount of data that is required to train regular neural ODEs is not available in many medical/biotechnological applications. This decreases the predictive capabilities that can be expected from neural ODEs in these settings. Second, while neural ODEs have the advantage of being flexible, the parameters of the neural network are not biologically inspired, which hinders the applicability in the biotechnological/medical domain. However, we can still leverage the highly efficient adjoint state method that is used to train neural ODE in metabolic kinetic model training. Furthermore, by mitigating parameter fitting challenges caused by characteristics of biological systems [15–17], as well as solutions from the field of neural ODEs [18, 19, 25], we stabilize the training process for systems biology models.

To showcase the usefulness of this neural ODE inspired training framework, we trained a large collection of SBML models for different priors on the parameters. Many of these models were retrieved from a previous benchmark [15], while others were downloaded from [27]. This allowed for a principled analysis of the effect of priors in parameter spaces, as well as further analysis of parameter importance.While many models were successfully trained, some models with more complicated dynamics such as oscillations were not quite as successful. Further work could be performed to increase the success rate by filtering parameter sets using periodic information using the Jacobian matrix [10].

To further reinforce the message that neural differential equations are useful for parameterizing kinetic models in real-world applications, we sought to train a previously established glycolysis model to fit a glucose pulse dataset from a feast/famine experiment [26]. This demonstrated that even for large-scale kinetic models, training could be performed within a few hours. We note however, that in this work the stoichiometry and flux mechanisms were known in advance, which is not always the case in the biotechnological/medical domain. For example, while the core metabolism of yeast is well-understood, other pathways might have unknown kinetics. Further work that focuses on augmentation of the rigid structure of kinetic models with a neural network might reduce the chance of fitting problems [59]. We envision that the *jaxkineticmodel* package is a first step in efficiently hybridizing ODE models [59].

## Materials and methods

### Data

#### SBML models

To rigorously test properties of Neural ODEs, we generate time-series metabolomics datasets from SBML models. The advantages of using synthetic data are that we can: 1) compare learned parameters against true parameters; and 2) test the method against a large collection of biological systems. The twenty-six models were retrieved from a previously reported set of benchmark models [15] and the Biomodels database [27]. The properties of the models are summarized in Table 1.

**Table 1.** Overview of the SBML models used in this study. The collection of models was retrieved from Biomodels [27] and a previously reported collection of SBML benchmark model [15]. The number of parameters, number of species, simulated range, and data points are reported.

| Model | Parameters | Species | $t_{end}$ | Data points |
| --- | --- | --- | --- | --- |
| Garde et. al 2020 [28] | 6 | 6 | 6 | 60 |
| Smallbone et. al (2013) [29] | 10 | 2 | 10 | 20 |
| Bruno et. al (2016) [30] | 10 | 6 | 100 | 60 |
| Beer et. al (2014) [31] | 12 | 4 | 8000 | 40 |
| Patil et. al (2023) [32] | 13 | 12 | 200 | 60 |
| Palani et. al (2011) [33] | 15 | 5 | 2500 | 50 |
| Crauste et. al (2017) [34] | 16 | 5 | 15 | 60 |
| Sneyd et. al (2002) [35] | 16 | 6 | 2 | 60 |
| Becker et. al (2010) [36] | 17 | 6 | 15 | 60 |
| Brannmark et. al (2010) [37] | 18 | 9 | 50 | 90 |
| Ray et. al (2013) [38] | 20 | 6 | 25 | 60 |
| Elowitz et. al (2000) [39] | 22 | 8 | 30 | 80 |
| Borghans et. al (1997) [40] | 24 | 3 | 3 | 30 |
| Fiedler et. al (2016) [41] | 24 | 6 | 10 | 60 |
| Fujita et. al (2010) [42] | 26 | 9 | 2000 | 90 |
| Bertozzi et. al (2020) [43] | 36 | 3 | 200 | 30 |
| Hass et. al (2017) [44] | 37 | 9 | 100 | 90 |
| Raia et. al (2011) [45] | 45 | 14 | 150 | 140 |
| Smallbone et. al (2011) [46] | 52 | 6 | 10 | 60 |
| Weber et. al (2015) [47] | 53 | 7 | 10 | 70 |
| Zheng et. al (2012) [48] | 62 | 15 | 100 | 150 |
| Isensee et. al (2018) [49] | 63 | 25 | 100 | 250 |
| Chassagnole et. al (2002) [50] | 117 | 36 | 10 | 360 |
| Messiha et. al (2014) [51] | 192 | 28 | 10 | 280 |
| Mosbacher (2023) [52] | 280 | 95 | 10 | 980 |

#### Feast/famine cycle and steady-state data

Neural ODEs are applied to datasets of a feast/famine cycle that was previously reported [26]. Feast/famine experiments are a stimulus-response experiment that consists of a 20 second feeding phase, followed by 380 seconds without feed. Three datasets were available for fitting a glycolysis model.

### Details on Neural ODE implementation

#### Code availability and used software

A tool to directly simulate SBML models in *JAX* was implemented using *Sympy* [53], *Diffrax* [21], and *LibSBML* [23]. All implementations for the experiment are publicly available.

#### Specifics of the gradient descent algorithm

The AdaBelief optimizer was used during training [54]. As previous reports show that systems biology models are better fitted in a log-transformed parameter space [15, 55], we applied the log-transformation of parameters before passing it through AdaBelief. Training was further stabilized by clipping the gradient global norm to prevent the exploding gradient problem often encountered in neural ODEs and recurrent neural networks [25]. This requires setting a maximum global norm hyperparameter (ĝ), which was set to ĝ = 4.

#### Scaling of species in the loss function

Biological systems exhibit large-scale differences in metabolite concentrations. This could result in the loss function J (mean squared error) being dominated by large absolute errors, even though the relative error might be small. To mitigate this, we use a similar strategy as proposed in [18], but instead of scaling by the minimum and maximum, we scale by the average concentration per metabolite over time (⟨m*_observed_*⟩), to prevent division by zero in steady-state data. Then, we calculate the mean squared error over N observed data points (eq. 2).

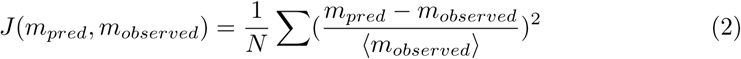

#### Specifics of the solver

Simulations were performed using the Kvaerno5 stiff ODE solver, which was already implemented in *Diffrax* [24]. The relative and absolute error tolerances were 10*^−^*^8^ and 10*^−^*^11^, respectively. The initial time step dt_0_ was set to 10*^−^*^10^. For the adjoint state method, the default implementation provided by *Diffrax* was used [21].

#### Gradient averaging for parameterization using multiple datasets

To showcase the practical usability of Neural ODEs, we aim to fit multiple datasets to the glycolysis model reported in the Supporting Information. These datasets are both steady-state metabolomics for different dilution rates, as well as dynamic data in the form of a glucose pulse. In order to simultaneously fit all datasets, we calculate the gradient dJ/dθ for each individual dataset and average them before updating using AdaBelief. This is a popular method to train a global model from local datasets in federated learning [60].

### Evaluating the training process on simulated data

Three properties of the training process are tested after training: 1) the initialization success rate of parameters; 2) the convergence properties through the relative improvement over the initialization; and 3) the distance of trained parameters to the true optimum. All experiments are performed three times.

#### Initialization success percentage

Datasets are simulated with the true parameters (θ*_true_*) that are specified in each SBML model. Latin hypercube sampling [56] is used to initialize 100 parameters within lower and upper bounds of the true parameter values, i.e., θ*_true_* /X ≤ θ*_true_* ≤ Xθ*_true_*. To investigate the effect of priors on initialization success, we chose priors for five different values of X: 2, 5, 10, 50, and 100. The initialization success rate, that is, the first iteration of stochastic gradient descent that leads to an estimate of the error, is calculated as a percentage of the total number of initializations.

#### Convergence of successful parameter initializations

For successful initializations, training is performed with 3000 iterations of stochastic gradient descent (AdaBelief) per initial parameter guess. To reduce computation time, training is stopped early when the loss function is below the loss threshold λ = 10*^−^*^6^. To quantify convergence properties, we look at the relative improvement in the mean squared error after training with respect to the initialization loss, as well as the percentage of successful training (λ < 0.001)

#### Global norm of gradients

The magnitude of the gradients with respect to the parameters is followed during the training process using the L2 norm (eq. 3). This allows to investigate the training process in more detail for different models.

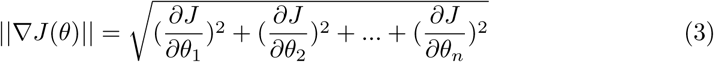

#### Comparing optimized parameters

To examine the uniqueness of parameter sets, log-transformed optimized parameters are compared to the true parameter values from which the synthetic dataset was generated using the cosine distance (eq. 4). This allows us to observe the variation in parameters. Optimized parameters are furthermore inspected using Principal Component Analysis (PCA).

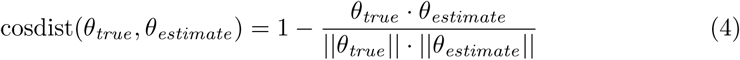

### Fitting the glycolysis model

To showcase the usefulness of the proposed framework we fit feast/famine datasets to a previously established kinetic model of glycolysis [57]. The model was reimplemented to be compatible with *JAX* to make parameters trainable [22]. The parameter initialization was done from parameters retrieved from literature. The model consists of 29 metabolites and 38 reactions, totaling 141 parameters. Twenty distinct rate laws were implemented as reusable *JAX* classes. Details on the implementation are reported in the supporting information (Table S1).

## Acknowledgments

The authors thank the AI4b.io consortium and Delft Bioinformatics Lab for their relevant discussions during progress meetings. We further specifically thank Liang Wu for his input on the final report. This work is supported by the AI4b.io program, a collaboration between TU Delft and dsm-firmenich, and is fully funded by dsm-firmenich and the RVO (Rijksdienst voor Ondernemend Nederland).

